# Evaluating a topic model approach for parsing microbiome data structure

**DOI:** 10.1101/176412

**Authors:** Stephen Woloszynek, Zhengqiao Zhao, Gideon Simpson, Michael P. O’Connor, Joshua Chang Mell, Gail L. Rosen

## Abstract

The increasing availability of microbiome survey data has led to the use of complex machine learning and statistical approaches to measure taxonomic diversity and extract relationships between taxa and their host or environment. However, many approaches inadequately account for the difficulties inherent to microbiome data. These difficulties include (1) insufficient sequencing depth resulting in sparse count data, (2) a large feature space relative to sample space, resulting in data prone to overfitting, (3) library size imbalance, requiring normalization strategies that lead to compositional artifacts, and (4) zero-inflation. Recent work has used probabilistic topics models to more appropriately model microbiome data, but a thorough inspection of just how well topic models capture underlying microbiome signal is lacking. Also, no work has determined whether library size or variance normalization improves model fitting. Here, we assessed a topic model approach on 16S rRNA gene survey data. Through simulation, we show, for small sample sizes, library-size or variance normalization is unnecessary prior to fitting the topic model. In addition, by exploiting topic-to-topic correlations, the topic model successfully captured dynamic time-series behavior of simulated taxonomic subcommunities. Lastly, when the topic model was applied to the David et al. time-series dataset, three distinct gut configurations emerged. However, unlike the David et al. approach, we characterized the events in terms of topics, which captured taxonomic co-occurrence, and posterior uncertainty, which facilitated the interpretation of how the taxonomic configurations evolved over time.

## INTRODUCTION

With the increasing availability of high throughput sequencing technologies, microbiome survey data is more readily available, which has allowed investigators to explore the use of complex machine learning and statistical methods to examine taxonomic diversity and extract relationships between taxa and samples. Nevertheless, many approaches struggle with the complexities inherent to microbiome data (1).

Microbiome abundance data are frequently generated via 16S rRNA marker gene surveys. This approach consists of sequencing the well-conserved 16S ribosomal rRNA gene from a set of samples, separating or clustering the resulting sequence reads into bins that capture taxonomic variation (e.g., Operational Taxonomic Units (OTUs), Ribosomal Sequence Variants), and then quantifying the proportion of these bins that originated from a given sample. The result is relative abundance data that is problematic for many analysis strategies.

For example, inadequate sequencing depth results in sparse count data, which in turn presents ordination artifacts for many dimensionality reduction techniques (2), renders many discrete linear regression models overdispersed (3), and biases estimates of microbial diversity and richness (3). Also, the dimensionality of the feature space relative to the number of samples makes microbiome data prone to overfitting, necessitating regularization (4). Yet another complication is that library sizes (the total number of sequence reads) differ, often considerably, between samples. Library size imbalance is a sequencing artifact and not representative of true biological variation. The consequence is that estimates of beta diversity is inflated due to undersampled taxa appearing rare (5). Thus, library size imbalance necessitates the use of relative rather than absolute abundances (6). These relative abundances are typically obtained by normalizing each count by its sample’s library size, but there are notable concerns with this approach. First, the degree of sparsity is increased by rounding small proportions to zero (3). Second, dividing each count by a common library size constrains each sample to the unit simplex (they must sum to 1), rendering many regression techniques and attempts to estimate covariance inappropriate (1,7). No longer can one effectively interpret regression coefficients; as one coefficient changes, the remaining coefficients must also change to satisfy the sum constraint. In other words, previously independent samples are now correlated due to their common denominator (8).

Strategies have been established to mitigate the obstacles that are unequal library sizes and “compositional” relative abundance data. A common approach, termed “rarefying,” involves down-sampling each sample’s library size to a common depth. The consequential loss of power can be drastic, however (9). McMurdie et al. (2014) proposed the use of variance stabilizing transformations on the raw count data for differential abundance analysis, thereby avoiding the sum constraint (9). Others have circumvented the sum constraint by using a centered log-ratio transformation (7,8,10) or isometric log-ratio transformations (6).

Beyond alternative normalization schemes, generative probabilistic approaches, such as Dirichlet-multinomial models (11–13), have garnered interest due to their appropriateness for microbiome data. Here, the microbial community is assumed to have been generated by a latent, community-related process. This interpretation is notably different from assuming that the sample represents overall community structure. Also, the use of a Dirichlet prior is appropriate given the discrete nature of microbiome data, and it too has a natural interpretation in this context; it can be viewed as the probability of sampling a specific microbial subcommunity (11). The relationship between Dirichlet and multinomial distributions have also laid the groundwork for probabilistic topic models, a dimensionality reduction technique that has been applied to microbiome data, modeling microbial source and sink environments (14), as well as inferring sample-taxa relationships (15). Despite their use, however, there has yet to be a thorough evaluation of the ability of topic models to capture underlying microbiome signal and whether normalization is necessary prior to model fitting

Here, we use a structural topic model (STM) to assess a topic model approach for 16S rRNA gene survey data, by evaluating its performance via simulation and via application to a well-known time-series dataset (16). In our first simulation, we show that library size normalization via rarefying or DESeq2 variance stabilization is unnecessary prior to fitting the STM, particularly for small sample sizes. Moreover, DESeq2 normalization results in a loss of power when identifying topic-sample effects. Our second simulation assessed the ability of topics to capture dynamic time-series behavior of taxa. We show that by exploiting topic-to-topic correlation, we can successfully recover predefined time-series interventions. When we applied our strategy to the David et al. (2014) dataset, a study that recorded the daily changes in microbiota for two individuals across two body sites (16), we recovered three distinct configurations of taxa that accurately represented the time-series events reported by David et al. However, unlike their approach, we characterized the events in terms of topics, which captured taxonomic co-occurrence, and posterior uncertainty, which facilitated the interpretation of how the taxonomic configurations evolved over time.

## METHODS

### Review of the Structural Topic Model

A topic model in the context of microbiome data is a Bayesian generative model that is fit to a vocabulary of N words (taxa) distributed across M documents (samples) (Table 1). The model aims to describe a sample as a mixture K latent topics (sets of co-occurring taxa), where each topic is described by a mixture of high frequency taxa. The model assumes that the probability of observing taxa xn in sample sm is given by 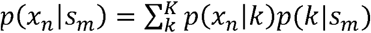 (17); thus, p(*x*_*n*_|*s*_*m*_) is influenced by the probability of observing x_n_ in topic k and the probability of observing topic k in sample s_m_. Biologically, topics consist of overlapping sets of co-occurring taxa that may share some biological context.

**Table 1.** Relationship of Terms

| Topic Model | Pipeline | Description |
| --- | --- | --- |
| Document | Sample, Day | Collection of reads from subject $m$ on day $d$ |
| Topic | Topic | Collection of co-occurring taxa, subcommunity |
| Word | OTU, RSV, Taxa | Features from taxonomic abundance table |
| Covariate | Sample feature | Sample-level variable of interest – e.g., event |
| $\theta$ (Sample-Over-Topics) | Topic frequencies | Frequencies of topics $\{1, \dots, K\}$ in sample $m$ |
| $\beta$ (Topics-Over-Words) | Taxa frequencies | Frequencies of taxa $\{x_1, \dots, x_n\}$ in topic $k$ |

The STM from (18) extends previous approaches such as latent Dirichlet allocation (19) by permitting the influence of sample covariates. The model is formulated as follows. Both a topics-over-taxa distribution β and a samples-over-topics distribution θ can receive sample information via their corresponding prior distributions. A logistic Normal (LN) prior is placed on β, where its mean is modeled as a linear combination of regression weights and sample covariates, in addition to a regularizing prior to prevent overfitting. Its covariance matrix allows for estimation of topic-topic correlations, providing a means to infer co-occurring topics over samples. θ, on the other hand, estimates the deviation of taxa frequencies from a background distribution (20). Word and topic assignments are generated via N- and K-multinomial distributions, respectively.

In the complete absence of sample covariates, the STM essentially reduces to a correlated topic model (21); with only θ prior covariates we have the Dirichlet-Multinomial regression topic model (22); and with only β prior covariates, we have a model analogous to a sparse additive generative model (20). Note that for all STMs described below, none will include sample covariates that influence the topics-over-taxa distribution β.

Posterior inference is performed via a partially semi-collapsed variational expectation maximization procedure.

### Simulation 1

Data were generated to assess (1) the ability of the STM to capture co-occurring sets of taxa, (2) the degree in which topics associate with sample covariates, and (3) the influence of library size imbalance and the need for normalization. S1 Fig shows our approach for simulation 1. We will refer to the figure sub-blocks 1-10 throughout this section.

(1a) To create the synthetic absolute abundance table (“balanced” table), we first generated a background distribution of size M(samples) × N(taxa) from a zero-inflated negative binomial distribution (ZINB) with sparsity (φ), mean (μ), and size (ψ) parameters adjusted to match distribution characteristics found in datasets such as Gevers et al. (e.g., sparsity, variance, max, etc.) (23). The ZINB was chosen given its ability to simulate the excessive zeros and overdispersion often encountered in 16S rRNA gene survey data (24). The dimensions of the taxonomic profile were a function of the number of samples M ⍰ {100, 500} and number of taxa N ⍰ {500, 1000} in a given simulation. (2) We randomly split simulated samples into equally sized treatment and control groups. (3) We then created 15 (arbitrary total) mock subcommunities (SCs) of size sc_l_ ⍰ {10, 15, 30}, composed of non-overlapping taxa that were generated by resampling with replacement all nonzero values in the background distribution and then scaling these values by effect size sc_m_ ⍰ {1, 2, 5, 10} and setting a proportion 1-sc_p_ (sc_p_ ⍰ {0.10, 0.25, 0.5,.0.75}) of these values to zero. Of these 15 SC_s_, 5 were set to replace the taxa abundances from a proportion g_p_ ⍰ {0.25, 0.50, 0.75} of treatment samples, 5 from a proportion of control samples, and 5 to replace an equal proportion from both treatment and control samples.

From the balanced table, we generated a second (“unbalanced”) table to investigate the effect of varying library size on model performance. (1b) Library sizes for each sample were randomly generated from a discrete uniform distribution [100, min(sample sum)] and used to resample the background distribution. (1c,d) The unbalanced table was then either rarefied to a balanced library size (N_min_=1000) (“rarefied” table) or normalized using the DESeq2 variance stabilizing transformation (“DESeq2” table) to create two relative abundance tables. Rarefying is a normalization approach where samples are down-sampled to a minimum value N_min_, and any samples falling below this value are discarded. While this approach does correct for library size biases, it has been shown to decrease statistical power (McMurdie & Holmes, 2014), which we hypothesized would have negative consequences when attempting to infer topic-sample covariate relationships. DESeq2 normalization is a variance stabilizing technique that adjusts discrete abundance data in terms of its mean-variance relationship and within-sample geometric mean. Recent work has shown it to be superior to rarefying (McMurdie & Holmes, 2014).

(4) After generating the rarefied and DESeq2 tables, STMs were fit to obtain thematic representations of the four simulated abundance tables. Model performance was assessed in two ways. First, we performed linear regression for each topic, using the frequency of topic k across samples (θ_,k_) as a dependent variable and the binary indicator for treatment and control as the independent variable. We will refer to the estimated regression coefficients as “topic-effects.” For each coefficient, we calculated 95% uncertainty intervals. Intervals that do not span 0 will be referred to as “detectable effects.”

(5-7) Second, we calculated Kullback-Leibler divergence (KLD) between p(x_n_|SC_w_)_data_ and p(x_n_|SC_w_,k)_model_, resulting in a distance for each topic-SC pair for a given model parameterization. It should be noted that while we chose KLD, we did explore the use of other information metrics such as Jensen Shannon distance; the results were analogous. (8-9) For a given STM with K topics, we identified the minimum threshold *th* in which there remain K KLD values less than *th* (equation 1):

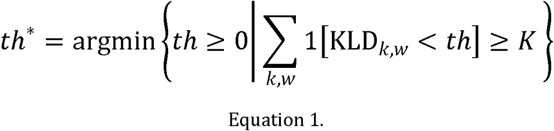

where K is the number of topics, KLD_*k,w*_ is the KLD between between p(x_n_|SC_w_,k)_model_ and p(x_n_|SC_w_)_data_ and 1 is the indicator function that returns 1 if KLD_*k,w*_ *< th* and 0 otherwise.

We posited that an outcome with good predictive power occurs when at least K topics mapped to a SC; fewer than K mapped topics guaranteed that some SCs were unaccounted for. These K values represent the K topics with smallest KLD to an SC. (10) We summed the number of SCs to which each of these K topics mapped (“redundancy scores”). Topics with small KLD to multiple SCs (a large redundancy score) would imply an inability of the topic model to separate SCs and thus capture their unique co-occurrence profiles. An ideal result would be each topic mapping uniquely to a single SC. We consider a many-to-one mapping acceptable, where multiple topics map to a single SC, as long as the topics map to one and only one SC.

#### Assessing simulation 1 performance

To infer the relationship between simulation parameters and threshold value, we performed multiple regression with the following scaled and centered covariates: number of taxa in a SC, number of total taxa, number of total samples, proportion of samples receiving the SC, SC effect size, SC sparsity, number of topics, and normalization method. For the normalization factor (DESeq2, rarefied, unbalanced, balanced), we set “balanced” as the reference level (i.e., the intercept). Threshold values were log transformed and used as the dependent variable. For redundancy score, we performed overdispersed binomial regression using the same set of covariates and setting K as the number of Bernoulli trials.

To assess the degree in which a given normalization procedure dampens topic-effects, for all parameter combinations, we quantified the proportion of detectable effects (topic-effects whose 95% uncertainty intervals did not span 0). We then performed overdispersed binomial regression with the following scaled and centered covariates: number of taxa in a SC, number of total taxa, number of total samples, proportion of samples receiving the SC, SC effect size, SC sparsity, number of topics, an indicator value representing whether a binary covariate for treatment verses control was present in the θ prior, and normalization method.

Quality of fit for all regression models was assessed by testing for equal variance and normality of the model residuals. Coefficients were considered statistically significant at p < 0.05.

#### Simulation 2

Synthetic abundance tables were created to assess the ability of the STM to detect time-series interventions that affect subsets of co-occurring taxa. Our approach was heavily influenced by the simulation detailed by Hall et al., who utilized Ananke to perform temporal clustering (25), but differs in the way we generated our synthetic abundance tables and our interventions. S2 Fig shows our approach for simulation 2. We will refer to the figure sub-blocks 1-3 throughout this section.

(1) We first generated 12 background distributions of 250 taxonomic features across 100 time points using the same ZINB distribution described in simulation 1. Then, we defined a SC as a set of 8 (arbitrary total) taxa. (2-3) We agitated various SCs by multiplying the background distribution by one of 3 types of interventions: pulses (S2 Fig: I1, I2), steps (I3, I4), and periodicity (I5, I6).

A pulse P is defined as a short term event where there is a mean shift in the background distribution for fewer than 5 time points T:

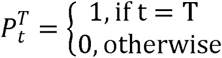

A step S extends from the initial intervention time point T_1_ until the end of the time-eries:

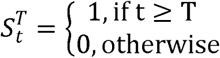

Periodicity P is defined as cyclical behavior that may or may not occur for the entirety of the time-series:

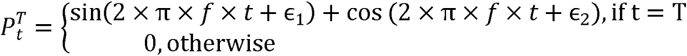

where f is the frequency of the signal and ε are the phase shifts.

Pulses and steps may include a weight that influences the rate of decay; that is, the rate in which the SC returns to its pre-intervention behavior (I2, I4). Using our set of interventions, we generated 12 synthetic time-series abundance tables. Samples were regarded as daily observations. All periodic interventions were fast, having a weekly period of 7 days (f=1/7). We posited that weekly periodicity is relevant to simulating gut microbiota dynamics.

For STM fitting, we treated the time index as a covariate representing day and created a second covariate representing day-of-week (DOW). Each synthetic dataset was fit with an STM (*K* ∈ {10,20,35,50,65,80,100}) that included a smoothing spline with 10 degrees of freedom on day and a second degree polynomial on DOW.

#### Event detection

For a given STM parameterization, we calculated the topic-topic correlation graph via Zhao and Lui (26), which is available in the R package stm (27) and wrapped in our package themetagenomics (28). Briefly, the procedure first performs a non-paranormal transformation on θ to alleviate the normality assumption. It then estimates the graph via the Meinshausen-Buhlmann method, which uses L1 regularization and is hence suitable for high-dimensional data (29). Selection of the regularization parameter was performed via the stability approach to regularization selection. We parsed the resulting correlation graph to identify cycles, linear chains, and clusters, which are defined as follows: a cycle consists exclusively of vertices of degree 2, forming a closed chain; a linear chain consists exclusively of vertices of degree 2, except at its ends, where each end may connect to a larger subgraph; and a cluster is a set of interconnected vertices of varying degree that may be connected to other subgraphs via a linear chain. The resulting subgraphs were used to identify correlated set topics that demonstrate similar behavior over time.

For each sample d, we generated 1000 θ_*d*,_ distributions from the posterior. Each posterior sample represents the topic frequencies at day d. With each θd, we sampled a topic assignment,followed by a taxa assignment from two multinomial distributions. For each topic assignment, we recorded its corresponding topic cluster defined by the topic-topic correlation graph.

#### Assessing STM performance

For each topic cluster from each of the 12 synthetic time-series, using the posterior predictive distribution for each STM fit, we calculated 4 statistics: cluster purity, cluster F1 score, cluster root mean square error (RMSE), and taxa RMSE. Cluster F1 score is a weighted average of cluster recall and precision. For each cluster c, we calculated the number of topic assignments belonging to cluster c that were sampled on days in which SC_w_ was present (true positives), the number of times topic assignments from cluster c were sampled on days in which SC_w_ was not present (false positives), and the number of times topic assignments from cluster c were not sampled on days in which SC_w_ was present (false negatives). F1 score is then defined as F1=2×TP/(2×TP+FP+TN).

For a SC of interest w, cluster purity represents the proportion of taxa x sampled from a topic belonging to cluster c that are members of SC_w_: p(x_n_ ∈SC_w_c). Purity was averaged over 25 sample batches to assess uncertainty.

Cluster RMSE was calculated as

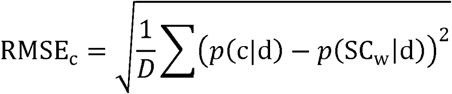

where *p*(c|d) is the frequency of cluster c being sampled on day d, averaged across 25 batches; p(SC_w_cd)is the frequency of SC_w_ on day d in the raw relative abundance table; and D is the length of the time-series. Taxa RMSE was calculated as

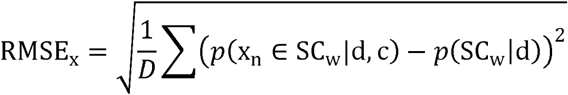

where p(x_n_ ∈ SC_w_d,c) is the frequency of the taxa sampled from cluster c belonging to SC_w_ and cluster c being sampled on day d.

#### Hierarchical clustering

We performed hierarchical clustering on the 12 synthetic time-series. This provided results from an alternative approach that we could use to further evaluate the STM performance.

For each synthetic time-series, we centered and scaled each taxon feature using the following equation: 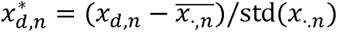. Then, for each time-series, we calculated 12 Euclidean distance matrices. Scaling each taxonomic feature enabled us to interpret the distances between features as a measurement of the differences in the shape of the signal, as opposed to differences in amplitude. Note that the library sizes across samples in this simulation were balanced, per the design of the simulation, hence no library size normalization was necessary. Hierarchical clustering was applied to each distance matrix using Ward’s minimum distance method. The resulting 12 trees were then cut to produce 30 HC clusters. The choice of 30 clusters was based on each SC containing 8 of the 250 total taxa for a given time-series. Because we are basing our choice for the number of clusters on what can be considered the true SC size, this can be considered a best-case-scenario. Performance was evaluated in terms of purity (defined above) and HC RMSE:

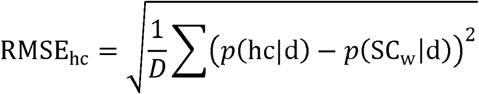

Number of clusters is the only free parameter for hierarchical clustering. We explored using DBSCAN as an additional clustering approach, but because it has two free parameters (number of clusters and signal radius), we felt hierarchical clustering was a more straightforward comparison.

#### Exploring Thematic Structure in David et al. 2014

##### Data Preparation and OTU Picking

The David et al. dataset contains fecal and salivary 16S rRNA surveys from two subjects. The samples were obtained at multiple time points across 318 days. Data from were downloaded from the European Bioinformatics Institute (EBI) European Nucleotide Archive (ENA) (accession number ERP006059). It consisted of 1.7 million 16S rRNA gene (V4 region) sequencing reads, 100 bp in length. The reads were quality filtered using the fastqFilter command in the dada2 package (30) with the following settings: trimLeft=10, truncLen=100, manN=0, maxEE=2, and truncQ=2. Closed reference OTU picking was then performed with QIIME version 1.9.1. using SortMeRNA again GreenGenes v13.5 at 97% sequence identity (31).

##### Data Preprocessing and STM Fitting

From the OTU table, we removed any samples with fewer than 1000 total reads, were not of fecal origin, were not from donor B, and did not include sample data for day, donor, and body site. OTUs lacking a known phylum classification or present in fewer than 1% of the remaining samples were removed. The remaining OTUs were normalized in terms of 16S rRNA gene copy number per the table provided by PICRUSt (32). The final OTU table consisted of 1562 OTUs across 189 samples.

We fit 7 STMs that varied in terms of topic number K ⍰ {15, 25, 50, 75, 105, 155, 250}. To infer the relationship between sample data and the samples-over-topics distribution θ, we used two sample covariates: two continuous, integer valued sequences representing days and DOW. Given our assumption that fluctuations in microbiota likely varied nonlinearly with respect to day, we used a smoothing spline with 10 degrees of freedom on day and a second degree polynomial on DOW.

##### Event detection

To detect events in subject B, we repeated the approach described for simulation 2. We compared our results to the 3 profiles described by David et al., which consisted of a pre-food-poising presentation (days 1-150), food-poising presentation (151-159), and post-food-poisoning presentation (150-318) profile.

##### Hierarchical clustering

We performed hierarchical clustering for comparison. The David et al. data were first centered log-ratio transformed to correct for library size imbalance. Each feature was then centered and scaled as described for simulation 2. Clustering was performed as detailed for simulation 2. The resulting tree was cut to produce 6 HC clusters. The choice of six clusters was based on the three profiles identified by David et al. (days 1-150, 151-159, and 160-318). We included three additional clusters to account for the background taxonomic variation lacking one of the three profiles of interest. Because we are basing our parameter choice on what can be considered the truth, this can be considered a best-case-scenario.

##### Measuring event effect size

We quantified the community-wide shift of taxonomic abundances with canonical correspondence analysis (CCA) and PERMANOVA (via the adonis function in the R package vegan (33)). For each synthetic time-series from simulation 2, we used binary indicators for each intervention as covariates. We then calculated the proportion of constrained inertia and R^^2^^ for CCA and PERMANOVA, respectively. We repeated this approach using the David et al. dataset, using two covariates, where covariate 1 was 1 for days 1-150 and 0 otherwise, and covariate 2 was 1 for days 160-189 and 0 otherwise.

## RESULTS AND DISCUSSION

Assessing the quality of topics is difficult. While we could compare the topics obtained by the STM to the sets of co-occurring taxa obtained by other methods, we would still be unable to verify whether the topics are biologically meaningful. Confirming co-occurrence via laboratory experiments would be ideal, but unrealistic in most circumstances. Thus, we turned to simulation where we were able to define the ground-truth – sets of taxa that co-occur across multiple samples (termed "subcommunities" (SCs)), which we hypothesize should be recoverable as topics. We had four objectives: (1) determine whether library size normalization improved the recoverability of co-occurring sets of taxa, (2) infer how library size normalization influences power for topic-sample covariate effects, (3) evaluate how robust the STM is to fitting complex, correlated signals that span across multiple samples, and (4) devise a topic model approach to capture complex signals of this type.

In simulation 1, we evaluated the influence of library size normalization on the ability of the structural topic model (STM) to (1) capture co-occurring taxa and (2) detect topic-sample covariate effects. Three approaches were compared to synthetic data with a balanced library size: down-sampling via rarefying, variance stabilization via DESeq2, and no normalization. In simulation 2, we evaluated the ability of the STM to capture simulated time-series events (termed “interventions”), that affected predefined SCs. Here, we leveraged correlated topics as a means of capturing topic dynamics over time. We compared our approach to the results obtained via hierarchical clustering. Lastly, we implemented our approach on time-series gut microbiome data from David et al. We focused on subject B, who notably presented with food poisoning midway through the study. We interpret the results in terms of topics and posterior uncertainty and compared our findings to those obtained by a hierarchical clustering approach, as well as the results reported by David et al. 2014.

### Effect of normalization on topic configurations (simulation 1)

#### For small sample sizes, unnormalized abundances as STM input data resulted in superior SC-to-topic mappings

We evaluated the ability of a given topic model parameterization to capture taxa co-occurrence by recovering predefined SCs. We found that, for small sample sizes (N=100), there was superior correspondence between SCs and topics when we used unnormalized abundances opposed to rarefied or DESeq2 normalized abundances. To quantify the effect normalization strategy had on the threshold value (our measurement of SC-to-topic correspondence), we performed multiple linear regression. With other covariates held fixed, relative to the balanced dataset, the threshold value was roughly twice as large for rarefied (β=0.397, SE=0.0309, p<0.0001) and DESeq2 (β=0.369, SE=0.0309, p<0.0001) normalized data compared to unnormalized data (β=0.189, SE=0.0309, p < 0.0001, R^^2^^=0.736). This indicated that both rarefying and DESeq2 normalization negatively affected the ability of topics to recover predefined SCs. This trend persisted irrespective of SC effect size and the number of taxa in a SC.

As sparsity decreased or sample size increased (N=500), the differences between normalization methods become less pronounced (Fig 1). This was largely due to the effect rarefying and DESeq2 normalization had on rare taxa. Rarefying down-samples taxa abundances; thus, rarer taxa are increasingly likely to not be resampled. DESeq2 normalization, on the other hand, can result in negative values for rare taxa. These values must be set to zero prior to STM fitting. Thus, in both cases, rare taxa have little to no influence on topic estimation, which likely impacted the ability of topics to map accurately to the predefined SCs.

**Fig 1.**
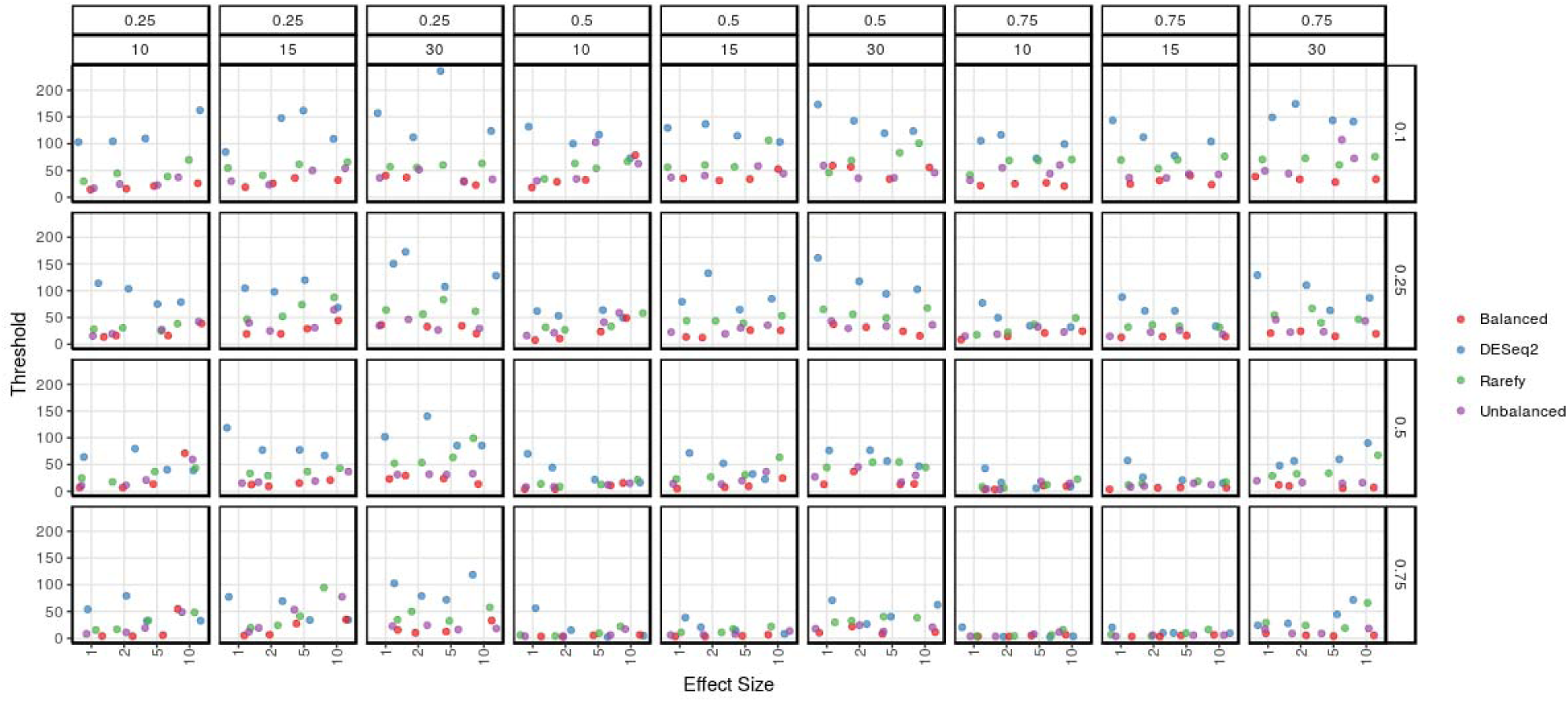
Simulation 1 (K25) threshold scores as a function of SC effect size. Simulated data consisted of 100 samples across 500 taxonomic features. Panel rows are ordered in terms of decreasing sparsity (1-scp); hence, large effect sizes for the bottom row equates to the largest SC signal. Panel columns are arranged by the proportion of samples containing the SC (top) and the number of taxa in the SC (bottom). Points are jittered and colored based on normalization method, where “balanced” indicated the simulated absolute abundances and “unbalanced” are the abundances after resampling with respect to library size. Small threshold values imply high correspondence between p(x_n_|SC_w_)_data_ and p(x_n_|SC_w_,k)_model_.

When we evaluated the number of topics with redundant SC mappings (topics that mapped to more than one SC), we found no relationship between redundancy score and either rarefied (β=0.022, SE=0.015, p=0.145) or unnormalized (β=0.019, SE=0.015, p=0.221) data, but found a positive association with DESeq2 normalized data (β=0.038, SE=0.015, p=0.013). This perhaps suggests that the dampening effect on rare taxa was greater for DESeq2 normalization than rarefying (to N_min_=1000), resulting in inferior topic mappings for DESeq2.

#### DESeq2 normalization is more conservative at detecting binary topic-sample effects

We next assessed the effect of normalization on detecting topic-effects. The number of detectable effects increased with increasing SC effect size scm and decreasing sparsity 1-scp (S3 Fig). DESeq2 normalization was the most conservative, frequently resulting in fewer detectable effects compared to balanced data (β=-0.169, SE=0.034, p<0.0001). In addition, it was the most sensitive to the presence of covariate prior information, increasing its detectable effects in 13/32 different combinations of effect size and sparsity parameterizations. Rarefying also negatively affected power, albeit less so compared to DESeq2 (β=-0.169 SE=0.034). Performing no normalization had little effect on the ability to detect topic effects (β=-0.059, SE=0.034, p=0.077). Increasing the total number of topics drastically diminished power for all normalization procedures (β=--0.593, SE=0.024, p<0.0001), particularly when the sample size was small (N=100). Of note, for K50, balanced data resulted in at most one detectable effect irrespective of parameterization. Increasing the sample size, however, resulted in a considerable increase in the number of detectable effects, with DESeq2 again behaving most conservatively.

Together, these results suggest that correcting for library size via DESeq2 normalization or rarefying is unnecessary, and possibly detrimental for small sample sizes. DESeq2 normalization decreased power for detecting topic-sample-covariate effects and slightly increased the frequency of redundant topic mapping relative to using balanced data. For small sample sizes, rarefying and DESeq2 normalization negatively affected the ability of the STM to recover SCs compared to using unbalanced abundances.

The performance of rarefying would likely improve with increasing Nmin, as shown in (34); however, many datasets are often under-sampled, necessitating the use of a small Nmin. The poor performance of DESeq2 normalization, on the other hand, is likely due to rare taxa receiving negative normalized values, which must be set to 0 prior to fitting the STM. This dampens the effect rare species have on inferring topic structure. A seemingly obvious adjustment would involve shifting the normalized values by a constant, but this is incorrect because the normalized values are in log-space (5). An alternative approach worth exploring could involve a centered log-ratio transformation using a Box-Cox transformation as opposed to a log transformation. While negative values would still occur, with the appropriate parameters, there may potentially be fewer, resulting in greater influence by rare species for topic estimation. Still, like DESeq2, this approach would require one to calculate the geometric mean across samples, which tend to be sparse. Thus, there is still need to identify an improved strategy for handling zeros when calculating the geometric mean, since using pseudocounts by simply adding a constant has been shown to yield spurious results (5,6)

#### Ability of topics to capture dynamic shifts in the configuration of taxa (simulation 2)

For the remaining sections, we will refer to distinct configurations of taxa spanning multiple time points as “profiles.” We will qualify this term accordingly: profiles identified in David et al. are terms “David profiles,” whereas those captured by the STM or hierarchal clustering are referred to as “topic profiles” and “HC profiles,” respectively. Contrast our use of “profile” with “cluster,” which we reserve for correlated topics found in the STM correlation graph (“topic clusters”) and clusters identified by hierarchical clustering (“HC clusters”). Multiple clusters in combination can together capture a particular profile.

#### Clusters of correlated topics successfully captured short-lived intervention dynamics

We evaluated the STM’s performance at capturing the behavior of multiple SCs across 12 synthetic time-series. We used four quality scores: F1, purity, cluster RMSE, and taxa RMSE. Fig 2 shows the scores for the best performing topic clusters for each time-series and SC (in terms of F1 score). The STM effectively recovered short-lived interventions (pulses) (sim 1; sim 3, SC 1; sim 5, SCs 2, 3, 4; sim 9, SC 1; sim 11; sim 12). The ten best scores for F1, cluster RMSE, and taxa RMSE all belonged to pulse interventions except the 9th largest F1 score (sim 6, k=10, SC=5). In addition, these clusters mapped well to their corresponding SC’s taxa; the top ten clusters in terms of F1 score had purity scores ranging from 0.421 (error = +/-0.060) to 0.628 (+/-0.063), suggesting that roughly half of all taxa populating these clusters were SC members. When we ranked the sampling frequency of all 250 unique taxa sampled from these 10 clusters, no SC member ranked lower than 23rd.

**Fig 2.**
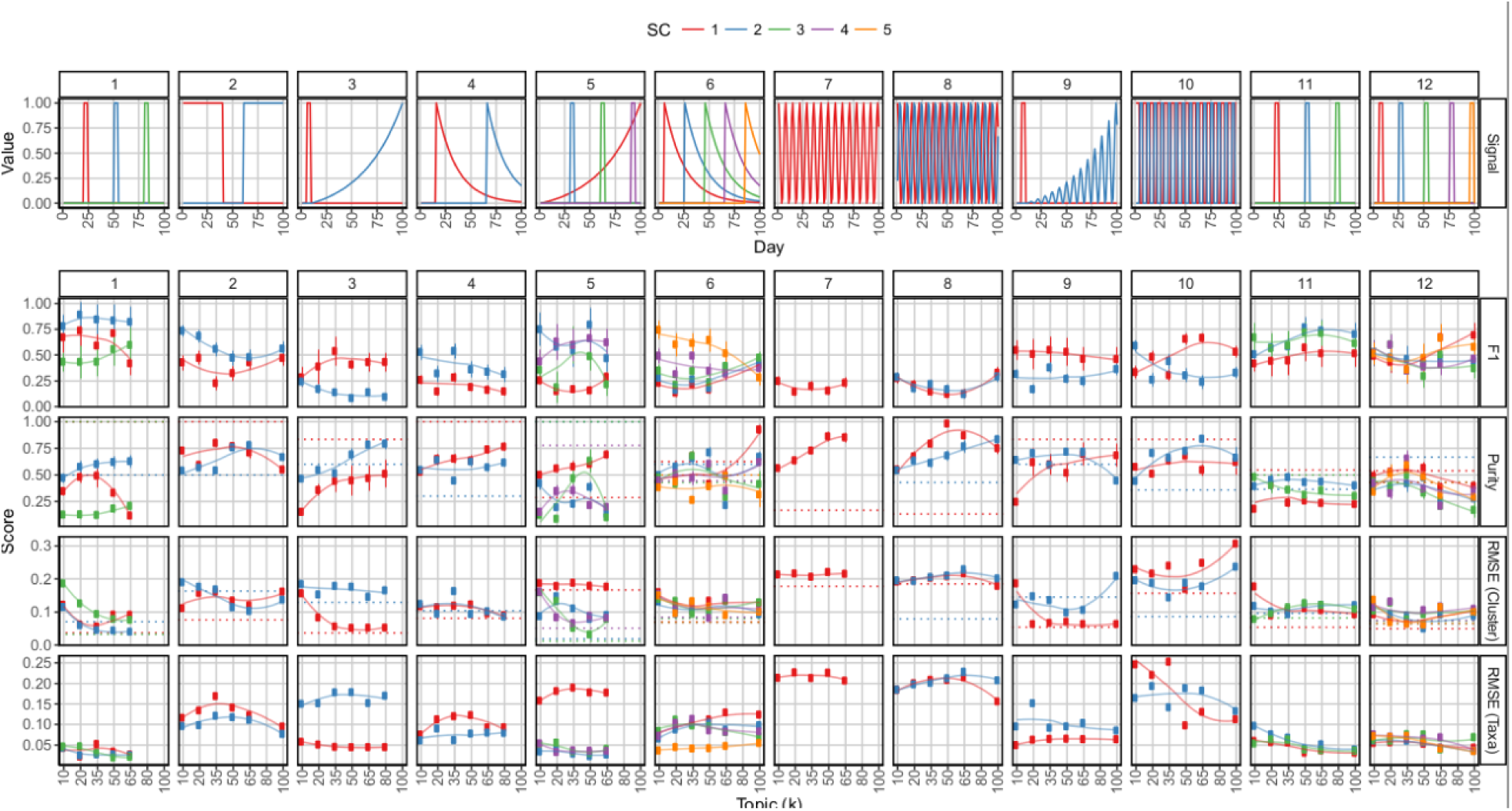
Simulation 2 interventions used to agitate the twelve 100 × 250 background distributions (top) and performance scores as a function of STM topic number (bottom). Panel rows contain the performance scores F1, purity, cluster RMSE and taxa RMSE. Panel columns contain the scores for each synthetic time-series after the corresponding interventions were applied. Colors correspond to a given SC. Hierarchical clustering (k=30) RMSE and purity scores from top performing clusters are shown as horizontal dotted lines.

The STM identified topic-SC mappings that had slightly worse RMSE compared to the RMSE for the HC clusters. For every time-series and SC, RMSE was lower roughly 19% of the time for the STM clusters compared to the HC clusters. There was a significant difference in mean RMSE (paired t-test, t=5.370, df=32, p<0.001), but not mean purity (t=-1.235, df=32, p<0.226). While this result suggests that hierarchical clustering outperformed the STM, note that we based the number of clusters (30) on our knowledge of how many taxa made up a SC (8) and how many taxa there were in total (250). Real-world datasets would lack this luxury. Moreover, because the choice of 30 clusters facilitated optimal HC cluster size, the resulting RMSE from the raw data would be at a minimum as long as the taxa making up the cluster well-approximated the true SC composition. Thus, the hierarchical clustering RMSE should be considered an ideal but improbable target.

#### Clusters of correlated topics recovered the periodic signals and outperforming hierarchical clustering in terms of purity

For the STM, the periodic signals (sims 7, 8, 10; sim 9, SC2) posed a difficult task because multiple topics tended to capture different segments of a long-term signal, making reconstruction of the signal difficult. Segmentation of a given signal was likely influenced by the sparsity-promoting priors, as well as increasing topic number.

Nevertheless, we hypothesized that the topics that captured neighboring segments of the complete time-series signal would likely be correlated across samples. This led us to parse the STM’s topic-topic correlation graph to identify subgraphs connected with non-zero edges, which we termed “topic clusters.” When visualized, it was apparent that the best performing clusters managed to capture the periodicity for each SC (Fig 3). Still, the performance of topic clusters in capturing long-term period behavior was worse compared to short-term interventions. Periodic signals resulted in poorer F1 scores and larger RMSE.

**Fig 3.**
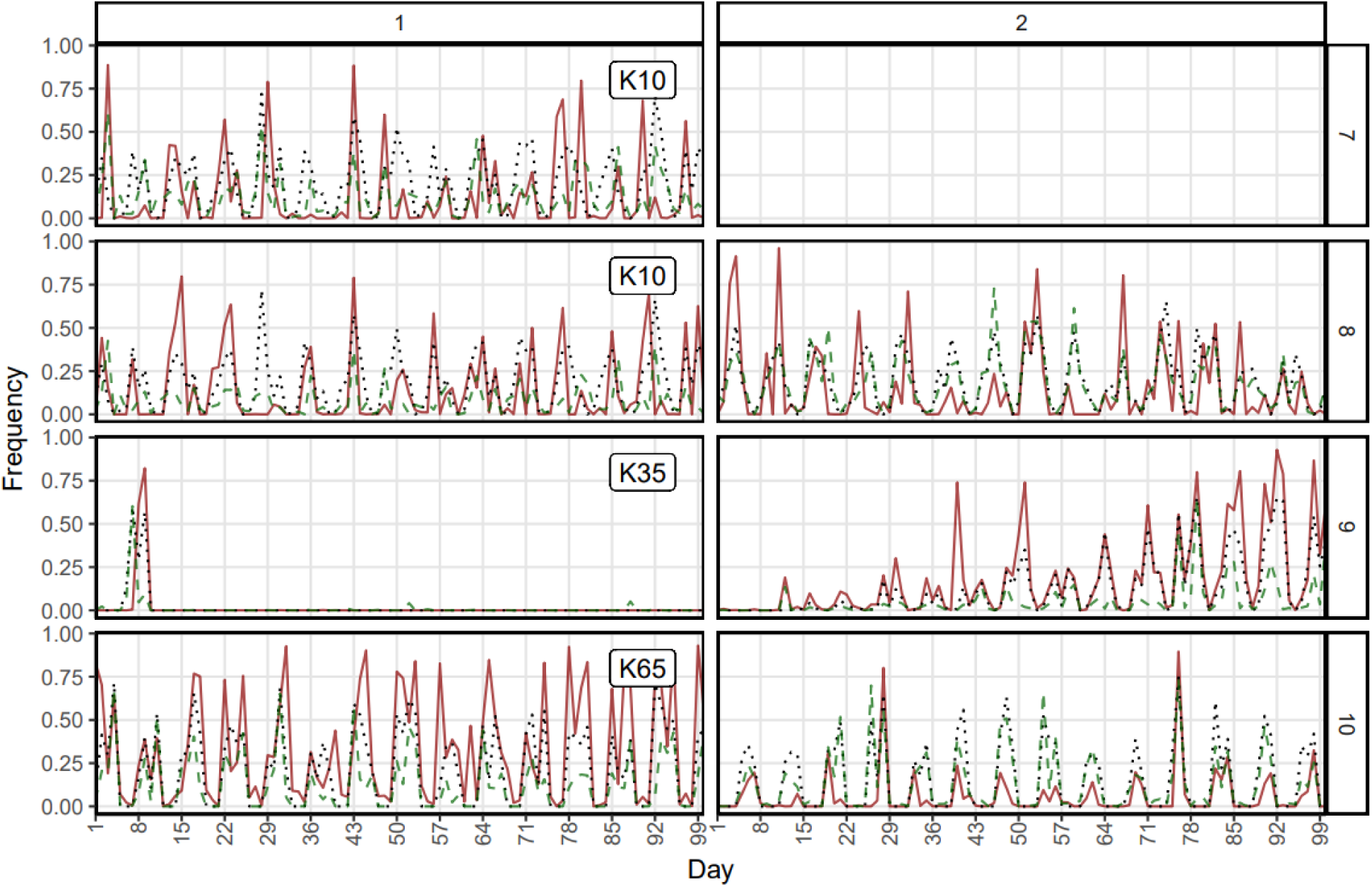
Mapping between p(SCperiodic|day) (black, dotted), p(HC*|day) (green, dashed), and p(cluster*|day) (red, solid), where * significant it is the best performing cluster in terms of F1 score or RMSE (for the STM and hierarchical clustering (k=30), respectively) for the SCs in a given simulation (row, 7-10). SCperiodic include only SCs that had periodic interventions (7-10). Columns show the SCs in a particular simulation. The topic number of the STM that yielded the best performing cluster is labeled in each row.

For periodic signals, the best performance was for SC1 in simulation 10, using 50 (F1=0.655 +/-0.054) and 65 topics (F1=0.666 +/-0.048). This simulation is notable for periodic signals that do not overlap, such that for a given week, SC1 spanned only the first 4 days, whereas SC2 spanned only the remaining 3. Simulation 8, on the other hand, involved two periodic SCs that were sinusoidal, with one SC phase-shifted. Interestingly, the STM managed to capture the taxonomic profile for each of the four periodic SCs. For K50 STMs, no top-performing cluster had a purity score of less than 0.679, with simulation 8, SC 8 performing best at 0.980 +/-0.014.

The STM clusters outperformed hierarchical clustering in terms of purity for periodic interventions: for 5/6 SCs, purity was larger for the STM clusters compared to the hierarchical clusters. Moreover, the average purity for the 6 HC clusters was 0.420, with two clusters as low was 0.167 and 0.133, suggesting an inability for hierarchical clustering to adequately capture the composition of periodic SCs. On the other hand, mean STM cluster purity was 0.761.

We also explored PCA as a means to reconstruct the time-series signals. S7 Fig shows the reconstructed signal for each of the 12 time-series, which suggests that PCA could capture the underlying signal. However, because we lacked a straightforward approach to recover the underlying taxa that compose a particular signal, we had no way of calculated RMSE to compare to the other approaches. This limitation alone suggests that using a PCA to capture dynamic SC behavior is limited.

#### Interventions with overlapping taxa negatively affected topic purity

Purity suffered the most for the time-series with overlapping SCs (sims 11-12), despite acceptable F1 scores and RMSEs. In simulation 12, K35 for SC 4 had the highest purity: 0.654 +/-0.076. Simulation 11 performed worse with a top purity score of 0.480 +/-0.039 (K10, SC 3). Roughly half of all clusters in simulations 11 and 12 had purity scores less than 0.388. The inability of topic clusters to adequately capture the SC profiles was due to topic clusters mapping to multiple SCs. For example, for cluster 7 in simulation 11, K10 mapped to SCs 2 and 3, which shared 4 taxa. This also suggested why taxa RMSE was lower than cluster RMSE. For a given posterior sample corresponding to day d, a topic cluster associated with multiple SCs may be drawn, negatively affecting cluster RMSE; however, only topics with high probability of being sampled at day d will be drawn, which in turn are likely to be linked with the SC associated with day d, positively affecting taxa RMSE.

In sum, exploiting topic-topic correlations provides a means to capture topic dynamics over time. Short lived dynamics are better captured by the STM; however, complex, long term behavior can be modeled, especially in circumstances where the complex signals do not overlap. Moreover, substantial mixing over OTUs may hinder interpretability in that topic clusters will correspond to multiple latent SCs. Still, one may still be able to separate overlapping SCs by manually parsing the individual topics that compose a correlated topic cluster.

### Detection of Events in Subject B from David et al

#### The STM identified 3 distinct gut configurations

In the topic correlation graph, we identified a cycle of three topics and two large subgraphs that contained 24 and 14 topics each (Fig 4A). The large subgraphs were connected by a linear chain of four topics (T9, T24, T2, T37). We defined the four sets of correlated topics as topic clusters and sampled, from the posterior, topic assignments and taxa assignments that fell into these clusters (Fig 4B).

**Fig 4.**
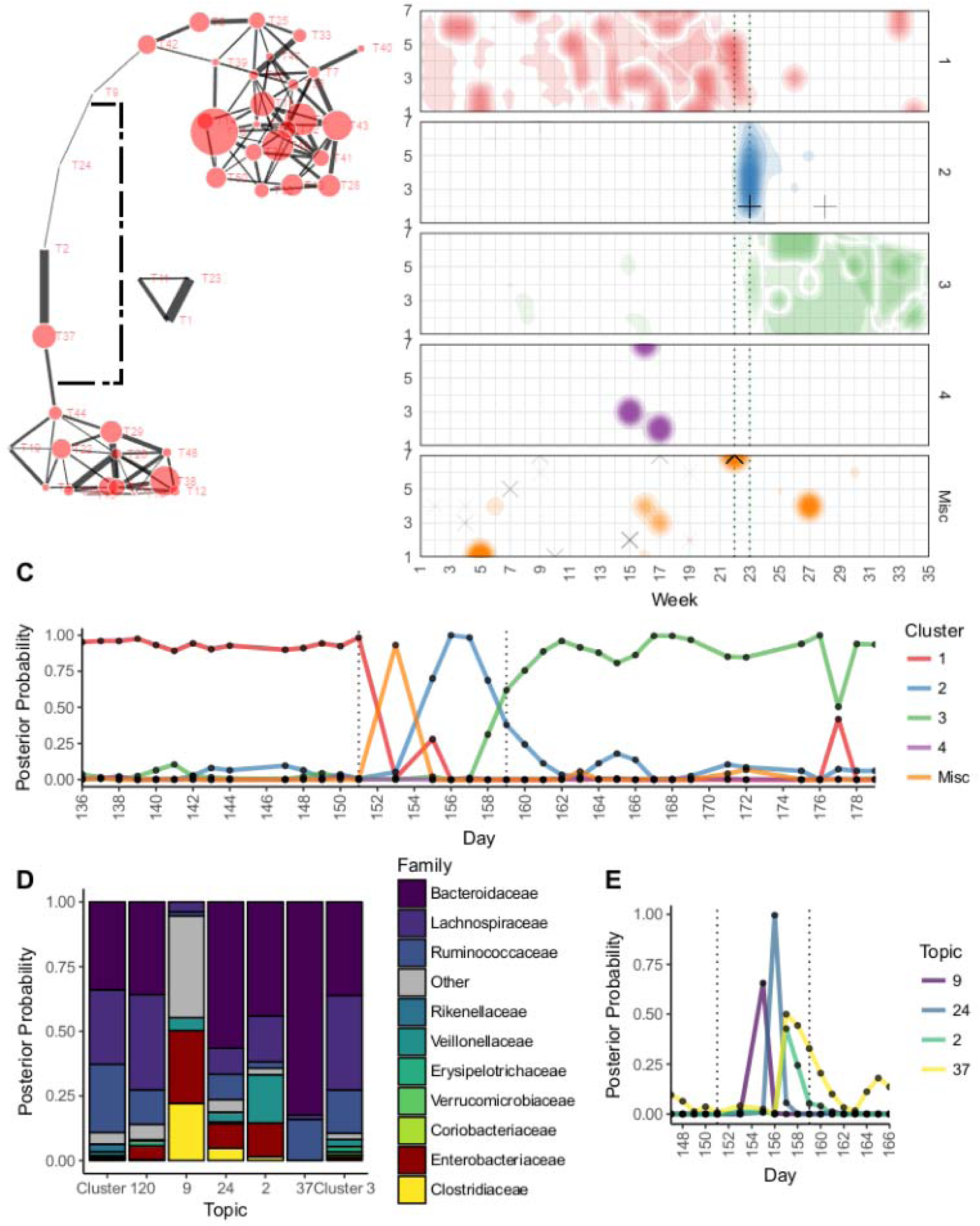
STM results for David et al. data. (A) The topic-topic correlation graph showing two topic clusters (clusters 1 and 3) connected by a linear chain (cluster 2). (B) Distribution of topic assignments as a function of day (week, DOW) and cluster (panels). The interval in which food poisoning symptoms presented (per David et al.) are marked with dotted vertical lines. Crosses indicate topic 9 assignments, whereas x’s mark topic 20 assignments. Uncertainty of topic assignments is expressed by the color transparency (more transparent implies greater uncertainty). (C) Frequency of cluster assignments as a function of day. (D) Frequency of taxa assignments given a cluster assignment. Cluster 2 is shown in terms of its topics (9, 24, 2, 37). Topic 20 is also shown (misc. cluster), which lacked any edges in the correlation graph, but marks the initial appearance of *Enterobacteriaceae*. (E) Frequency of topic assignments as a function of day for cluster 2. The shift in frequency mirrors its order in the correlation graph.

There were two clear delineations between the distribution of topic assignments for the 3 clusters, specifically when transitioning from cluster 1 to 2 (weeks 22-23; days 152-154) and clusters 2 to 3 (weeks 23-24; day 161). Our intervals are similar to the original study’s transition points at days 144-145 and 162-163, where the shift from a cluster 1 to cluster 2 profile corresponded with subject B’s food poisoning diagnosis.

Because we can assess the uncertainty in θ and hence the uncertainty in both topic and taxa assignments, we can characterize the shift in the gut profiles over time as a function of posterior probability (Fig 4C). The transition between clusters 1 and 2 is abrupt and likely occurred around day 153. Taxonomically, this transition is marked by a shift from Bacteroideaceae (posterior probability=0.338), Lachnospiraceara (0.276), and Rumunococcaceae (0.266) to Enterbacteriaceae (0.246) and Clostridiaceae (0.195) families (Fig 4D). In particular, day 153 was distinctive for topic 20. This rare topic was not correlated with any other topics and hence did not belong to any topic cluster. While its taxonomic profile was quite similar to cluster 1, it was distinctly enriched for *Enterobacteriaceaea spp*., which is consistent with the subject’s Salmonella diagnosis. Topic 20 likely marks the event of initial exposure to the pathogen.

The distribution of topic assignments for cluster 2 followed the order in which its topics were positioned in the topic correlation graph (the linear chain) (Fig 4E). The start of the cluster 2 profile, day 155, was dominated by topic 9, characterized by a profile substantially different from cluster 1. Bacteria enriched in this topic included *Haemophilus parainfluenzae, Clostridium perfringens*, and, notably, Enterobacteriaceaea spp. Thus, topic 9 likely represented the disrupted configuration of microbiota due to exposure to Salmonella. Enterbacteriaceae spp. and C. perfringens, via topic 24, continued to dominate on day 156. Day 157 was best described by topic 2, a topic rich in *Enterobacteriaceae spp*. as well as *Veillonella spp*. It should be noted, however, that our results were more conservative than David et al. in that we confidently estimated the cluster 2 profile lasted roughly 4 days (155 to 158), which is much shorter than the original study’s estimate (145 to 162). Our estimated length of illness (153 to 158) was more consistent to David et al. (151 to 159), however.

At approximately day 159, the gut profile shifted toward cluster 3, a profile similar to cluster 1 in terms of Bacteroidaceae (0.369), but enriched in Lachnospiraceae (0.360) and depleted in Rumunoicoccaceae (0.165) (Fig 4D).

#### Hierarchical clustering resulted in a wider estimate for the length of the illness profile

With hierarchical clustering, we created six clusters based the three profiles reported in David et al. (S8 Fig.) Note that we did explore other parameterizations, which yielded similar cluster configurations with respect to both time and taxonomic composition (S9-10 Figs.). Since we used a priori knowledge, identification of these clusters can therefore be considered a best-case-scenario. Three clusters (2, 3, 6) corresponded to the days in which subject B presented with food poisoning. Clusters 5 and 6 were comprised of 355 and 298 taxa, respectively, and, in the raw relative abundance table, both peaked on roughly days 151 to 157. However, the taxa in these clusters during this span were low-frequency taxa; all had mean relative abundance less than 0.0002. In cluster 5, the taxa with largest mean relative abundance included *H*. *parainfluenzae, Leuconostocaceae spp., Dialister spp., and Enterobacteriaceae spp*., whereas cluster 6 included *Klebsiella spp., Closridiaceae spp*. and *Enterobacteriaceae spp*. Cluster 2, on the other hand, spanned days 151 to 169, and contained taxa considerably larger in terms of mean relative abundance: *Bacteroides spp*. (mean relative abundance=0.192), *Enterbacteriaceae spp*. (0.034), and *H. parainfluenzae* (0.013) composed this cluster. Together, these three clusters likely correspond to profile 2 identified by David et al. (days 145 to 162) and clusters 5 and 6 (151 to 157) specifically correspond to the time of illness estimated by both the STM (153 to 158) and David et al. (151 to 159). However, unlike the STM approach, these clusters consist of substantially more taxa and hence are inundated with more noise.

Cluster 4 contained 360 taxa and corresponded well to the pre-illness period, spanning days 1 to 150. During this span, large mean relative abundance taxa that associated with cluster 4 included *Bacteroides spp* (0.156), *Lachnospiraceae spp.* (0.078), and *Faecalibacterium praunitzii* (0.050). This set of taxa was similar to the taxa identified in the STM’s profile 1. The post-illness period (profile 3) was captured by clusters 1 and 3, but these clusters failed to completely separate profile 2 from profile 3; they spanned days 151 to 318. They were composed of taxa similar to cluster 4, but with a substantial contribution from the family Ruminococcus, a change seen in profile 3 for the STM. The top mean relative abundance taxa in clusters 2 and 3 were *Ruminococcus spp*. (0.132), *F. prausnitzii* (0.112), *Bacteroides spp*. (0.086), and *Lachnospiraceae spp* (0.025).

These results suggest that the profiles identified in the STM are similar to those obtained via hierarchical clustering. However, the sparsity inducing priors in STM ease interpretation since the profiles are less contaminated with unimportant taxa. The smallest cluster obtained with hierarchical clustering contained 121 taxa (cluster 1). Without prior knowledge to suggest where the breaks between profiles may occur, identifying meaningful abundance profiles (during the tree cutting stage or the analysis stage) may be increasingly difficult. Also, the STM identified topics that likely represented the initial presentation of the illness (day 153, topic 9) and a sequence of topics that shows a gradual evolution of the abundance profile (topic cluster 2). The clusters associated with disease obtained via hierarchical clustering unsuccessfully separated the shift from profile 2 to 3 and, moreover, where unable to demonstrate how the profiles evolved over time.

#### Shifts in the taxonomic abundance profiles for the synthetic time-series from simulation 2 were similar to the shifts observed in the David et al. data

Given how clear the delineations between shifts in gut profiles were, we attempted to quantify the degree in which the David et al. profiles changed before and after the subject’s bout with food poisoning. Doing so enabled us to compare the signal seen in David et al. to our synthetic datasets from simulation 2. We used proportion of inertia (via CCA) and R2 (via PERMANOVA) for a given signal as our measure of effect size. The results are shown in S1 Table, which indicate that the David et al. signals represent slightly less total variation compared to the synthetic datasets, with the periodic datasets 7 and 8 being most similar.

## CONCLUSION

We have demonstrated a topic model approach for 16S rRNA gene survey data. By evaluating its performance via simulation, we have shown that it is unnecessary to perform library size normalization via rarefying or DESeq2 variance stabilization prior to fitting the STM. DESeq2 normalization results in a loss of power when identifying topic-sample effects, especially when sample size is small (N=100). We have also shown the ability of topics to capture dynamic time-series behavior of taxa. We exploited topic-topic correlation to successfully reconstruct predefined time-series interventions. Our approach was best at reconstructing short lived interventions. Despite worse performance when modeling periodic interventions, the STM outperformed a hierarchical clustering approach, with ideal parameters, in terms of purity. When we applied the STM approach to the subject B gut microbiome data from David et al. (2014), we recovered three distinct configurations of taxa that agreed with the results of David et al. Unlike their approach, however, we characterized the events in terms of topics, which captured taxonomic co-occurrence, and posterior uncertainty. This enabled us to describe the evolution of these taxonomic configurations over time. Compared to hierarchical clustering, the STM approach resulted in sparser taxonomic clusters, improving our ability to capture meaningful signal relative to noise. In addition, unlike hierarchical clustering, the STM successfully separated the transition between taxonomic profiles 2 and 3.

Future work should focus on methods capable of integrated the benefits of dimensionality reduction obtained using a topic model approach with sophisticated zero replacement and normalization techniques. While our results suggest that such transformations may be unnecessary, we contend that the poor performance of DESeq2 was largely due to dampening the influence of rare taxa when setting negative normalized values to zero. More appropriate strategies may overcome issues stemming from overdispersion and zero-inflation while mitigating the biases that result directly from normalization and zero replacement strategies.

Our topic model approach is available in our package themetagenomics (35).

---

**S1 Fig.**
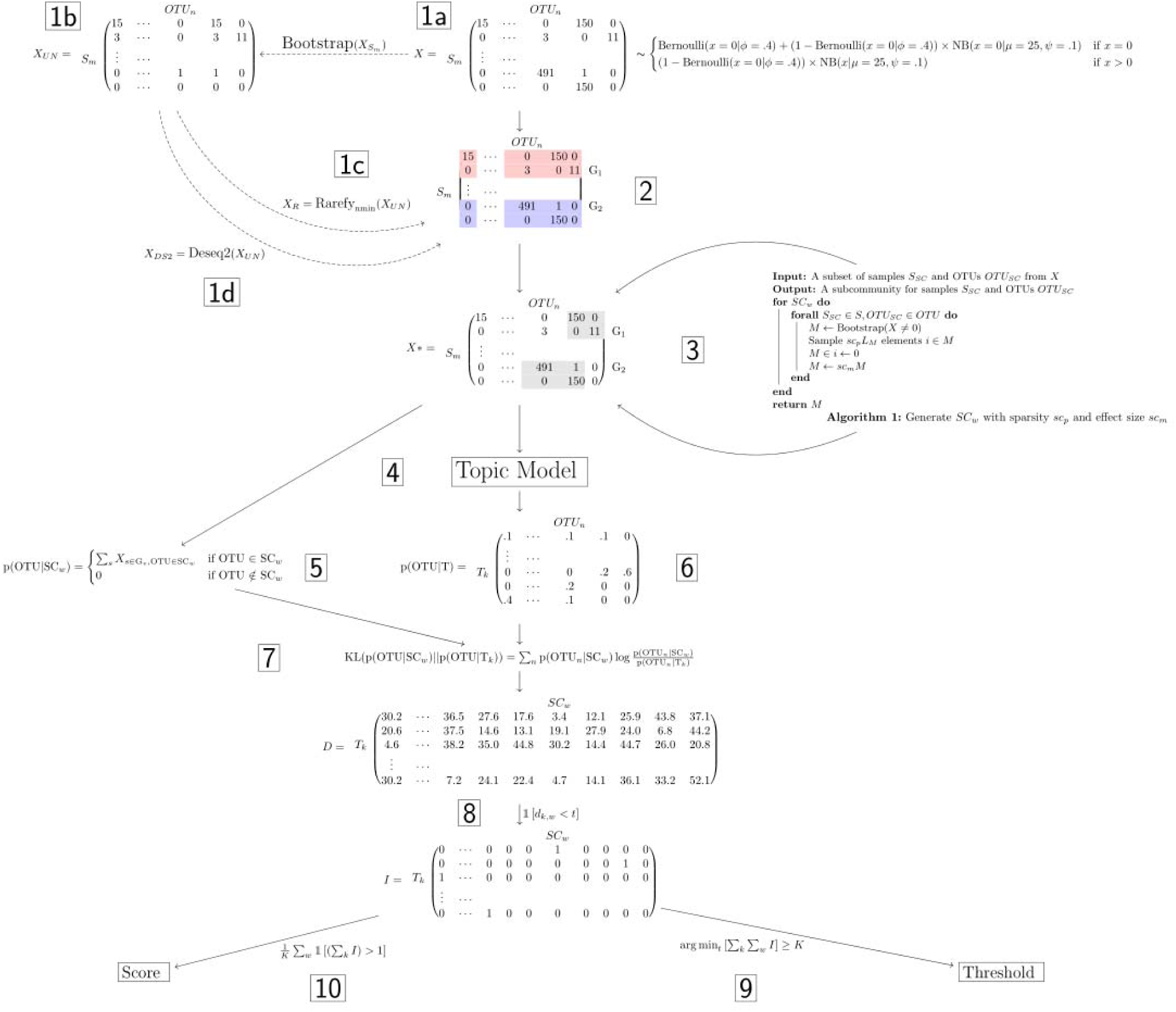
Workflow for simulation 1. (1a) A background distribution generated from a zero-inflated negative binomial distribution (ZINB) with sparsity (φ), mean (μ), and size (ψ) parameters. (2) Samples were randomly split into treatment (G1) and control (G2) groups. (3) 15 subcommunities (SCs) of size sc_l_ ⍰ (10, 15, 30) were generated by resampling with replacement all nonzero values in the background distribution and then scaling these values by effect size sc_m_ ⍰ (1, 2, 5, 10) and setting a proportion 1-sc_p_ (sc_p_ ⍰ (0.10, 0.25, 0.5,.0.75)) of these values to zero. 5 SCs each were set to replace the taxa abundances from a proportion g_p_ ⍰ (0.25, 0.50, 0.75) treatment samples, control samples, and an equal proportion from both treatment and control samples. (1b) Library sizes for each sample were randomly generated from a discrete uniform distribution [100, min(sample sum)] and used to resample the background distribution. (1c,d) This table was then either rarefied to a balanced library size (1000) or normalized using the DESeq2 variance stabilizing transformation to create the two additional abundance tables. (4) STMs were fit. (5-7) We calculated Kullback-Leibler divergence (KLD) between p(x_n_|SC_w_)_data_ and p(x_n_|SC_w_,k)_model_, resulting in a distance for each topic-SC pair for a given model parameterization. (8-9) For a given STM with K topics, we identified the minimum threshold th in which there remain K KLD values less than th. (10) We summed the number of SCs to which each of these K topics mapped (“redundancy scores”).

**S2 Fig.**
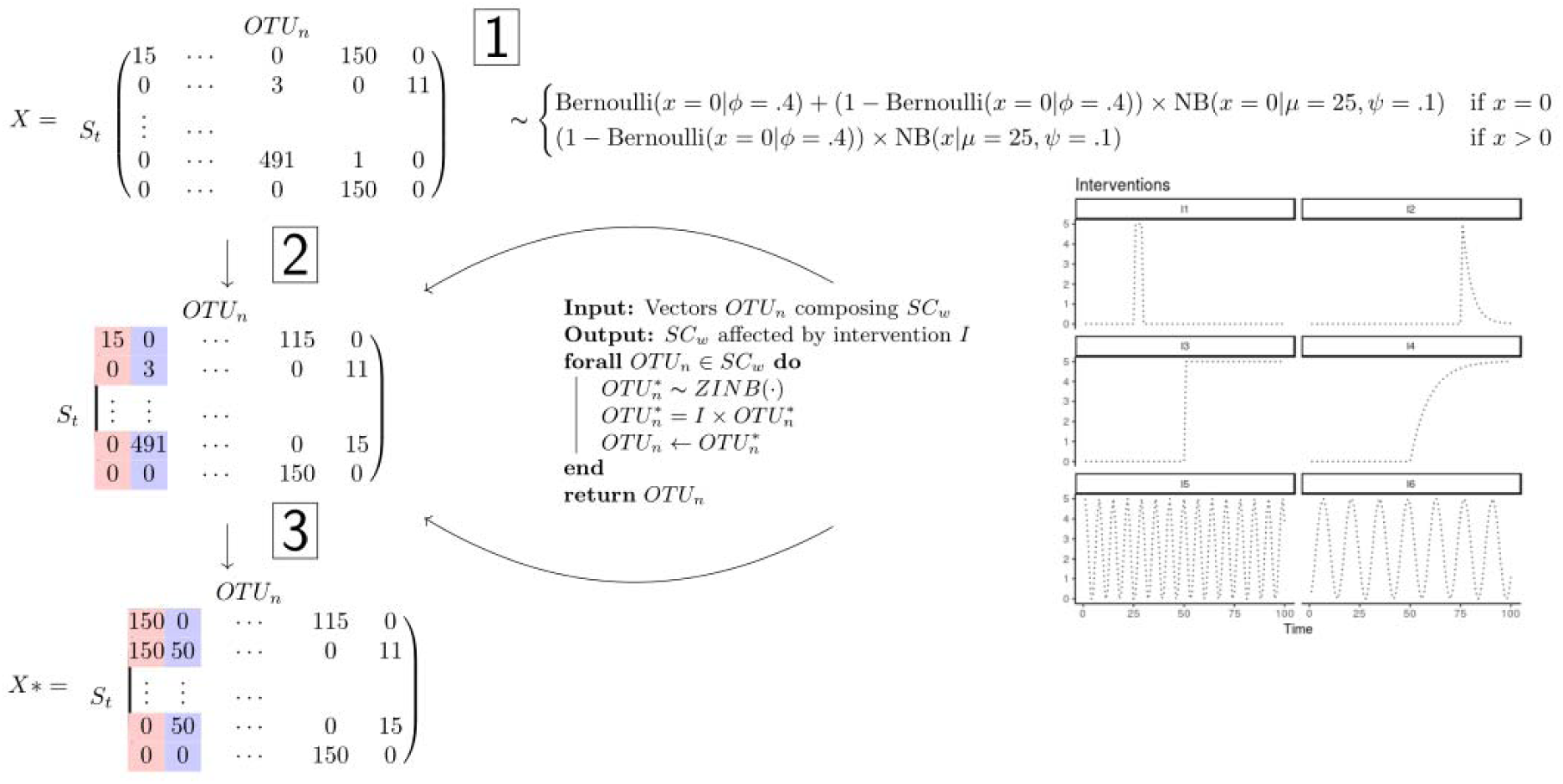
Workflow for simulation 2. (1) 12 background distributions were generated from a zero-inflated negative binomial distribution (ZINB) with sparsity (φ), mean (μ), and size (ψ) parameters. (2-3) Each SC of 8 taxa were agitated with one or more interventions: pulses (I1, I2), steps (I3, I4), or periodicity (I5, I6).

**S3 Fig.**
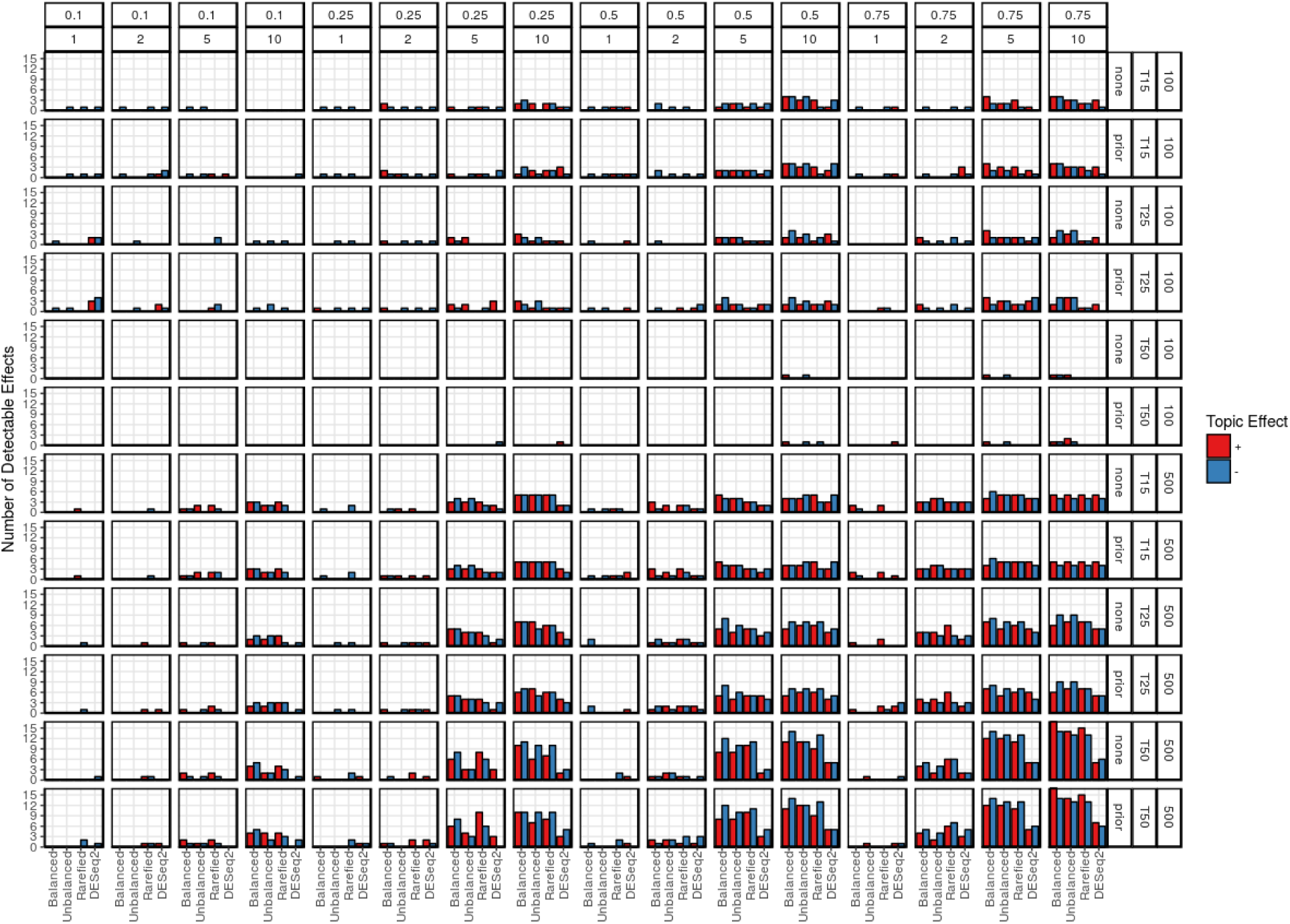
Simulation 1 detected effects as a function of normalization method. Panel rows are ordered in terms (1) presence of prior information (binary indicator for treatment group), (2) number of topics, and (3) sample size (100 samples, 500 taxonomic features; 500 samples, 1000 taxonomic features). Panel columns are arranged in terms of decreasing SC sparsity (1-scp) (top) and SC effect size (bottom). Bars are colored based on the direction of the detectable effects, where positive effects (associated with the treatment group) and negative effects are red and blue, respectively. We consider results for balanced data (absolute abundances) as a best-case-scenario; hence, significant deviations from the effects detected for balanced data would suggest poor performance in terms of type 1 or type 2 errors.

**S4 Fig.**
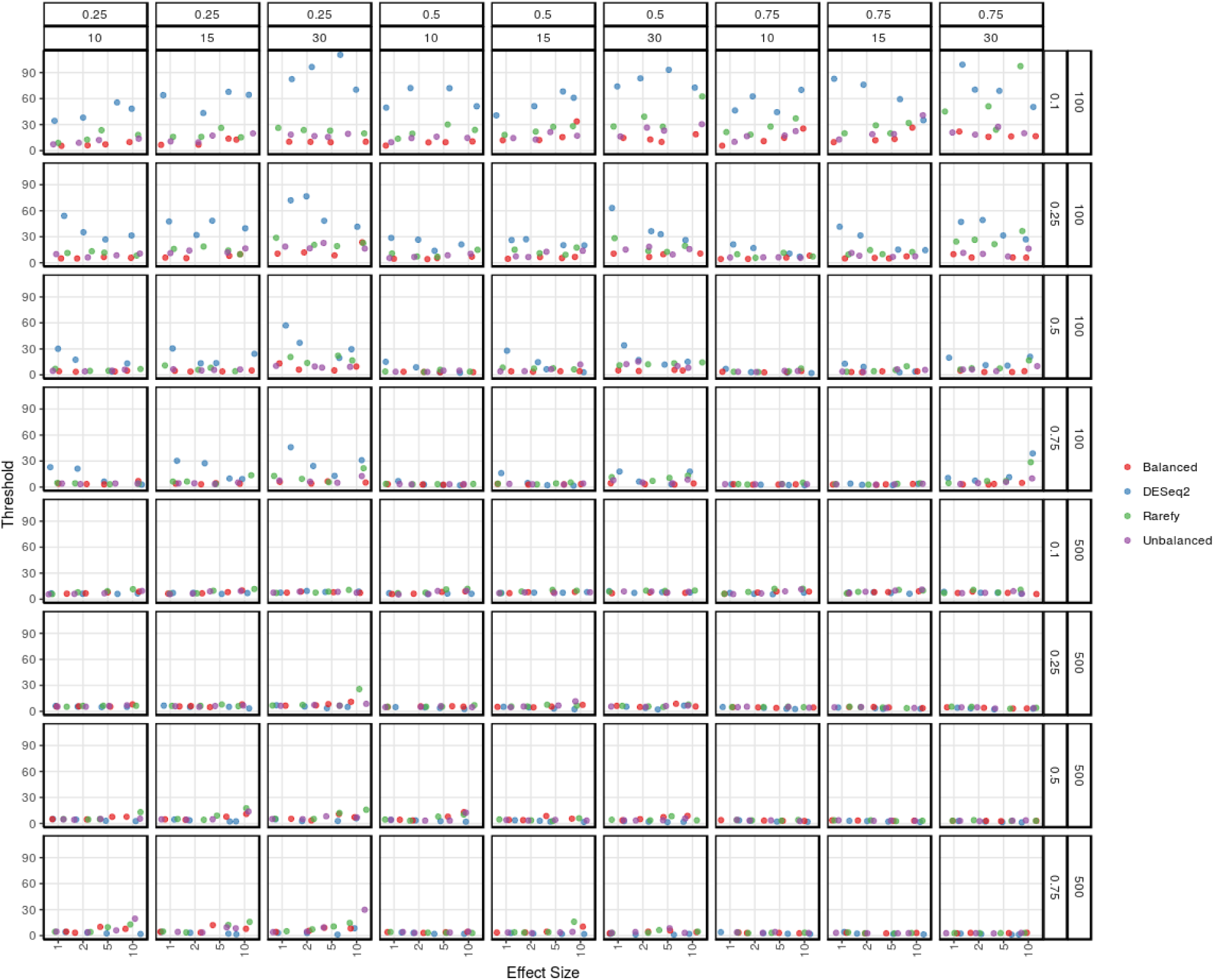
Simulation 1 (K15) threshold scores as a function of SC effect size. Panel rows are ordered in terms of decreasing sparsity (1-scp) and sample size (100 samples, 500 taxonomic features; 500 samples, 1000 taxonomic features). Panel columns are arranged by the proportion of samples containing the SC (top) and the number of taxa in the SC (bottom). Points are jittered and colored based on normalization method, where “balanced” indicated the simulated absolute abundances and “unbalanced” are the abundances after resampling with respect to library size. Small threshold values imply high correspondence between p(x_n_|SC_w_)_data_ and p(x_n_|SC_w_,k)_model_.

**S5 Fig.**
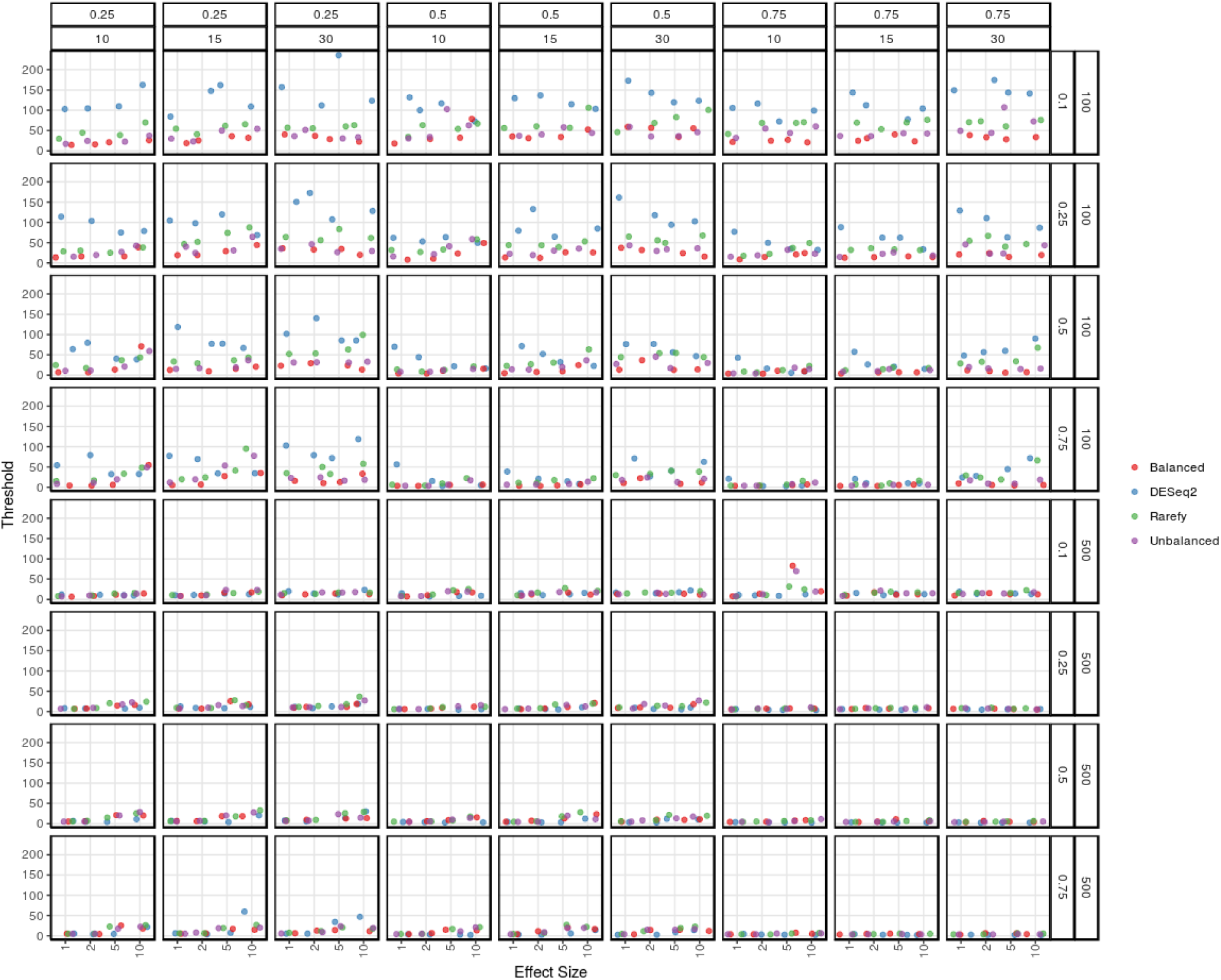
Simulation 1 (K25) threshold scores as a function of SC effect size. Panel rows are ordered in terms of decreasing sparsity (1-scp) and sample size (100 samples, 500 taxonomic features; 500 samples, 1000 taxonomic features). Panel columns are arranged by the proportion of samples containing the SC (top) and the number of taxa in the SC (bottom). Points are jittered and colored based on normalization method, where “balanced” indicated the simulated absolute abundances and “unbalanced” are the abundances after resampling with respect to library size. Small threshold values imply high correspondence between p(x_n_|SC_w_)_data_ and p(x_n_|SC_w_,k)_model_.

**S6 Fig.**
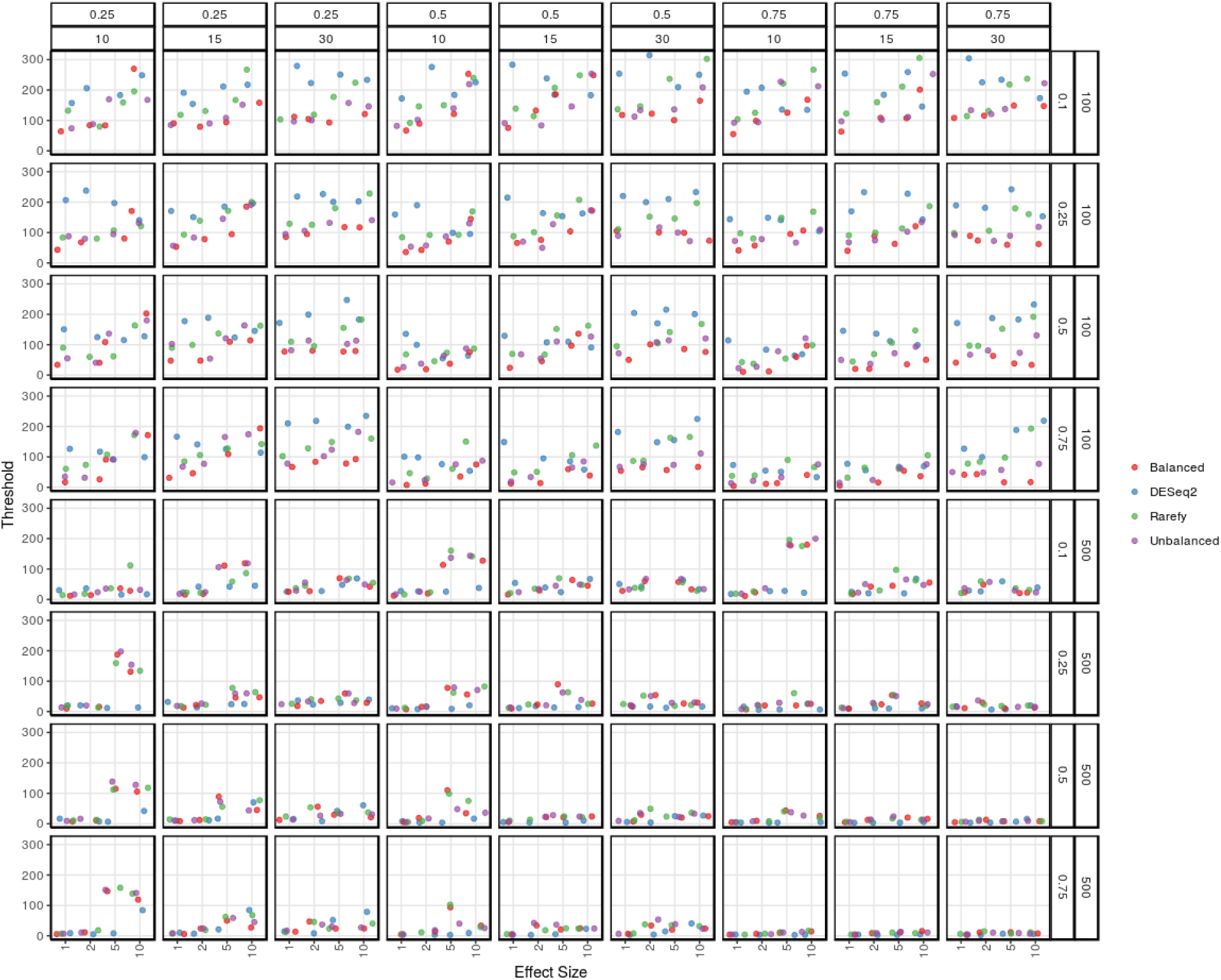
Simulation 1 (K50) threshold scores as a function of SC effect size. Panel rows are ordered in terms of decreasing sparsity (1-scp) and sample size (100 samples, 500 taxonomic features; 500 samples, 1000 taxonomic features). Panel columns are arranged by the proportion of samples containing the SC (top) and the number of taxa in the SC (bottom). Points are jittered and colored based on normalization method, where “balanced” indicated the simulated absolute abundances and “unbalanced” are the abundances after resampling with respect to library size. Small threshold values imply high correspondence between p(x_n_|SC_w_)_data_ and p(x_n_|SC_w_,k)_model_.

**S7 Fig.**
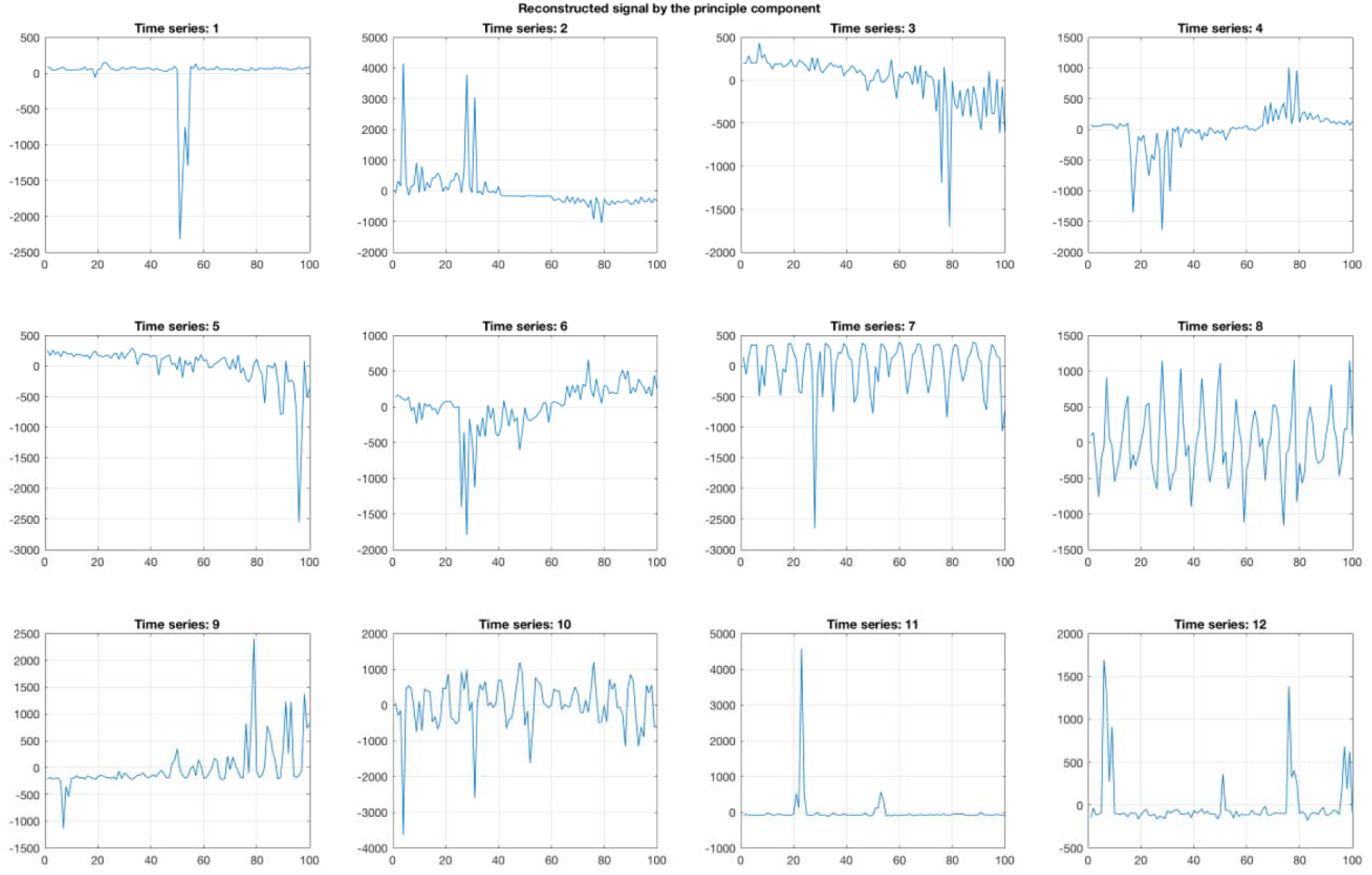
PCA reconstruction of David et al. data (subject B).

**S8 Fig.**
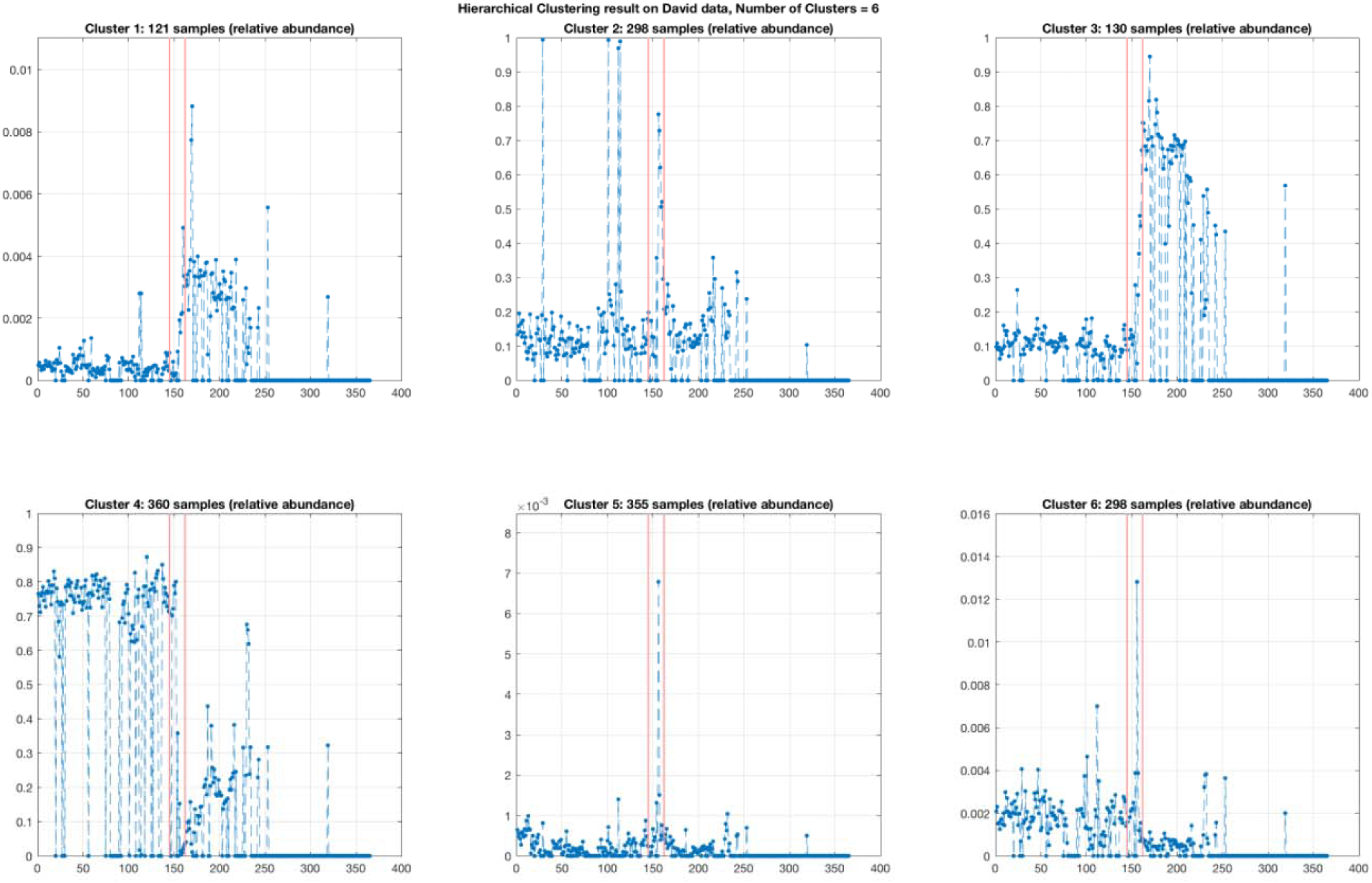
Clusters via hierarchical clustering (k=6) applied to the David et al. dataset (subset B). Red lines signify the presentation of illness.

**S9 Fig.**
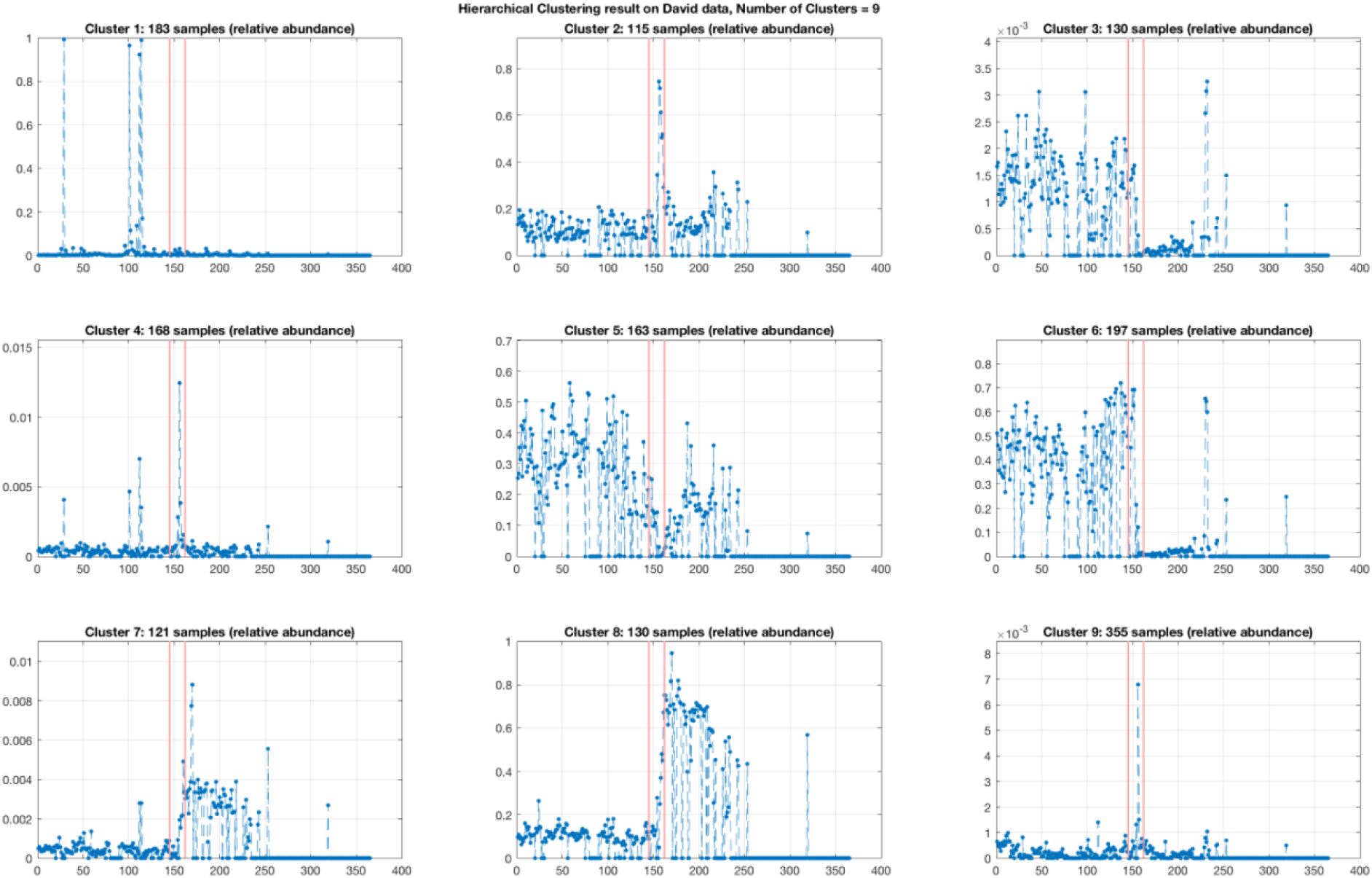
Clusters via hierarchical clustering (k=9) applied to the David et al. dataset (subset B). Red lines signify the presentation of illness.

**S10 Fig.**
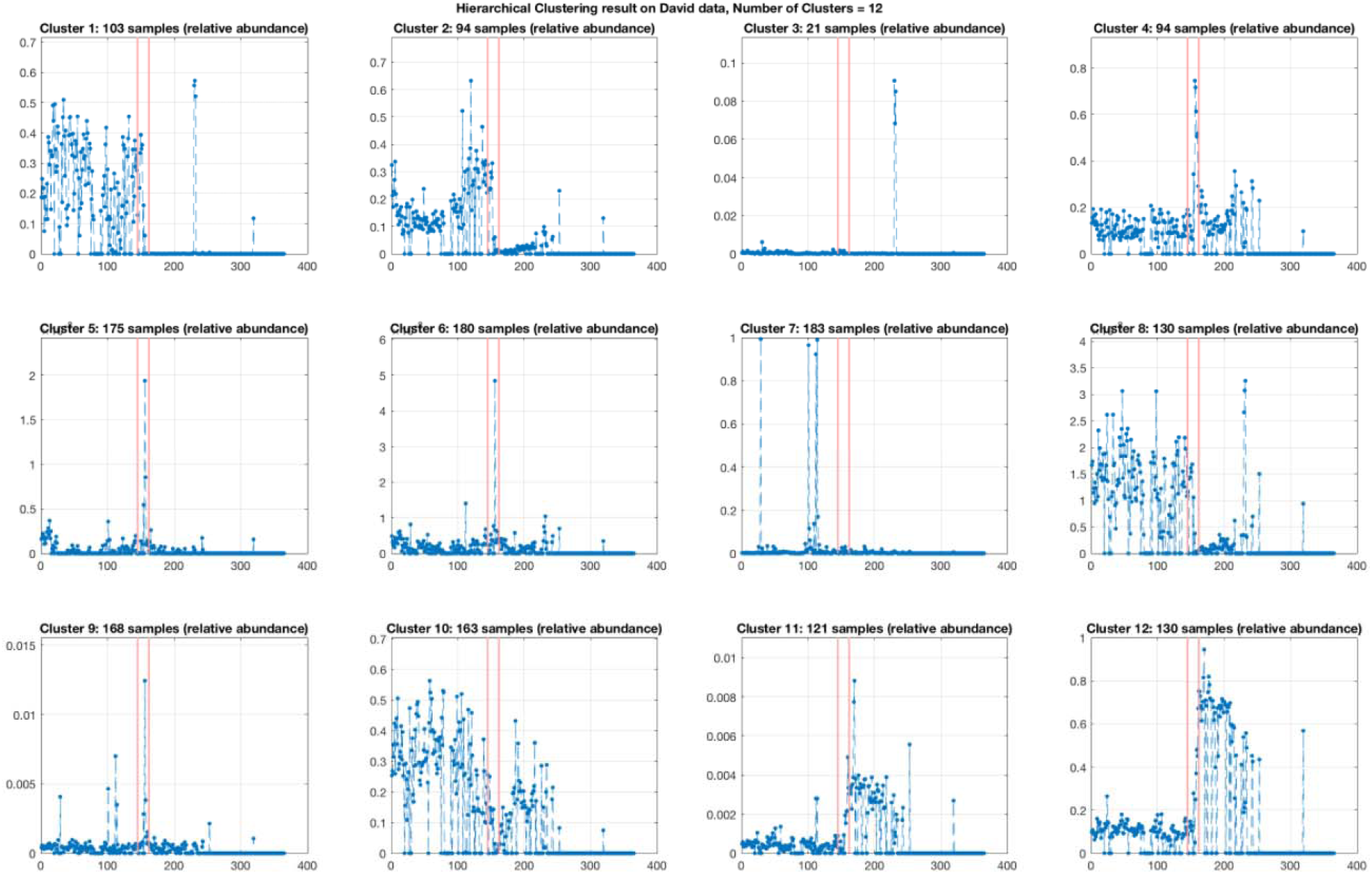
Clusters via hierarchical clustering (k=12) applied to the David et al. dataset (subset B). Red lines signify the presentation of illness.

**S1 Table.** Comparison of the measured time-series effect size between David et al. (D) and the simulations (1-12) from simulation 2. Inertia is the mean constrained inertia from CCA with the intervention(s) as a covariate. R^^2^^represents the variation explained by these covariates when performing PERMANOVA. Effect sizes closest to David et al. are shown in bold.

|  |  | Simulation |  |  |  |  |  |  |  |  |  |  |  |
| --- | --- | --- | --- | --- | --- | --- | --- | --- | --- | --- | --- | --- | --- |
|  | D | 1 | 2 | 3 | 4 | 5 | 6 | 7 | 8 | 9 | 10 | 11 | 12 |
| Inertia | **0.003** | 0.03 | 0.035 | 0.022 | 0.018 | 0.024 | 0.022 | **0.009** | **0.009** | 0.025 | 0.044 | 0.024 | 0.022 |
| R<sup>2</sup> | **0.008** | 0.060 | 0.087 | 0.033 | 0.066 | 0.067 | 0.179 | **0.012** | **0.021** | 0.032 | 0.139 | 0.061 | 0.098 |

